# A century of guayule: Comprehensive Genetic Characterization of the Guayule (*Parthenium argentatum* A. Gray) USDA Germplasm Collection

**DOI:** 10.1101/147256

**Authors:** Daniel C. Ilut, Paul L. Sanchez, Terry A. Coffelt, John M. Dyer, Matthew A. Jenks, Michael A. Gore

## Abstract

The fragility of a single-source, geographically concentrated supply of natural rubber, a critical material of the modern economy, has brought guayule (*Parthenium argentatum* A. Gray) to the forefront as an alternative source of natural rubber. The improvement of guayule for commercial-scale production has been limited by the lack of genomic tools and well-characterized genetic resources required for genomics-assisted breeding. To address this issue, we developed nearly 50,000 single nucleotide polymorphism (SNP) genetic markers and genotyped 69 accessions of guayule and its sister taxa mariola (*Parthenium incanum* Kunth), representing the entire available NALPGRU germplasm collection. We identified multiple interspecific hybrid accessions previously considered guayule, including six guayule-mariola hybrids and non-mariola interspecific hybrid accessions AZ-2 and AZ-3, two commonly used high-yielding cultivars. We dissected genetic diversity within the collection to identify a highly diverse subset of guayule accessions, and showed that wild guayule stands in Big Bend National Park, Texas, USA have the potential to provide hitherto untapped guayule genetic diversity. Together, these results provide the most thorough genetic characterization of guayule germplasm to date and lay the foundation for rapid genetic improvement of commercial guayule germplasm.

**Key Results:**

1. Six guayule accessions are guayule-mariola hybrids
2. Guayule collections from Big Bend National Park contain novel guayule genotypes not present in collections from Mexico
3. Commonly cultivated accessions AZ2 and AZ3 contain introgressions from other *Parthenium* species
4. The triploid accessions 11591, 11646, N576, N565, N565II, and RICHARDSON are generally indistinguishable from each other with respect to genetic background and likely represent the 4265-I source genotype (Johnson, 1950)
5. Open pollinated and purposefully outcrossed tetraploid selections derived from 4265-I incorporate further genetic diversity and form distinct genotypes

## Introduction

Natural rubber is a critical raw material of modern society, essential to a diverse range of industries such as automotive, electronics, clothing, and health care. However, natural rubber is currently a single-source material with geographically concentrated production. The rubber tree [*Hevea brasiliensis* (Willd. Ex A. Juss.) Müll. Arg.] provides over 99.9% of the world supply of natural rubber, and over 75% of it is produced in South-Eastern Asia (FAOSTAT; <u>http://faostat3.fao.org</u>). Plantations are generally clonal and vulnerable to the South American leaf blight fungal pathogen *Microcyclus ulei* (Rivano, 1997). In response to this situation, guayule (*Parthenium argentatum* A. Gray), a woody perennial shrub native to the desert regions of northern Mexico and southwestern United States, has been repeatedly assessed and utilized as an alternative source of natural rubber in the United States, and declared a critical agricultural material in times of crisis (7 U.S.C. § 171; 7 U.S.C. § 178).

The sporadic nature of research funding, germplasm collection, and stock maintenance (Hammond and Polhamus, 1965; Ilut et al., 2015, fig. 1; Thompson and Ray, 1989), combined with ploidy variation within guayule populations (Gore et al., 2011; Ilut et al., 2015) and the unusual guayule reproductive biology—sporophytic self-incompatibility in diploid plants and facultative apomixis (diplospory type) in polyploid plants—have made it difficult so far to achieve significant increases in rubber yield, a critical step on the path to developing guayule as an alternative commercial source of natural rubber. In order to address this issue with modern genomics-assisted breeding approaches, a comprehensive characterization of ploidy variation and genetic diversity within the NALPGRU germplasm collection is essential. A previous limited study identified only two primary genotypic groups within the collection (Ilut et al., 2015), stressing the need for greatly expanded sampling of the germplasm collection in order to genetically enrich the guayule germplasm pool.

**Figure 1.**
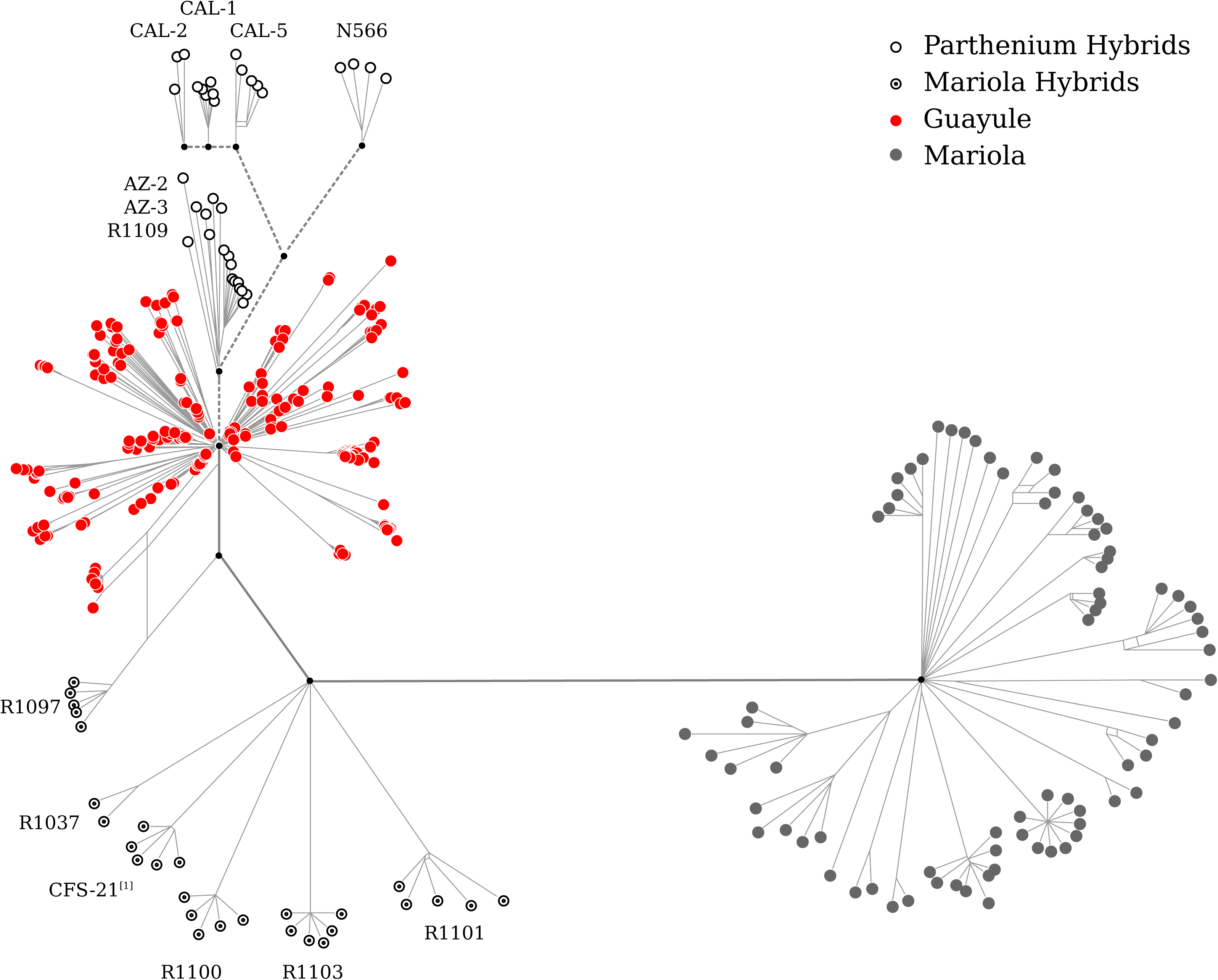
Phylogenetic relationships of accessions in the NALPGRU guayule and mariola germplasm collection. Terminal node colours and symbols indicate guayule accessions (red filled circles, center left), mariola accessions (gray filled circles, lower right), guayule-mariola hybrids (black open circle and dot, lower left), and other guayule hybrids (black open circle, top left). Hybrid accessions are identified with their respective accession names. Main branches connecting hybrids of guayule and unknown *Parthenium* species to the guayule clade are marked with dotted lines. Accessions containing a mixture of guayule, mariola, and guayule-mariola hybrids are denoted with the superscript [1].

We generated ploidy estimates and scored nearly 50,000 single nucleotide polymorphism (SNP) markers for 395 plants from 69 accessions of guayule, guayule hybrids, and mariola (*Parthenium incanum* Kunth), representing the complete collection of guayule and mariola plant material available in the NALPGRU germplasm repository. We identified six guayule-mariola hybrid accessions previously labelled as guayule, detected two high-yielding guayule cultivars as hybrids between guayule and an unknown *Parthenium* species distinct from mariola, and identified seven historic guayule accessions that contain broad mixtures of guayule genotypes. Finally, we selected two groups of accessions that represent distinct, diverse examples of guayule genetic diversity from central Mexico and southwestern United States, respectively, and provide a robust foundation for future exploration of genetic and phenotypic diversity within guayule.

## Results and discussion

### Sampling, genotyping, and SNP marker analysis

For each of the 69 accessions in this study, we sampled an average of six plants per accession and performed ploidy and genotyping-by-sequencing (GBS) analysis on each of those plants. Detailed results for each sample are available in Table A.1 (see Supplementary data) and accession-level summaries are presented in Table 1. For the purpose of phylogenetic analysis, the genotyping data were filtered to a high confidence subset of 48,495 SNP markers from which we identified four distinct clades (Figure 1) representing guayule germplasm, non-mariola *Parthenium* hybrids of guayule, guayule-mariola hybrids, and mariola germplasm. The SNP markers were further filtered within each clade to retain only those that were variable within the clade. A summary of these filtering results are presented in Table A.2, and detailed principal component analysis (PCA) results for all clades are available in Figures A.1-A.8 (see Supplementary data).

**Table 1.**
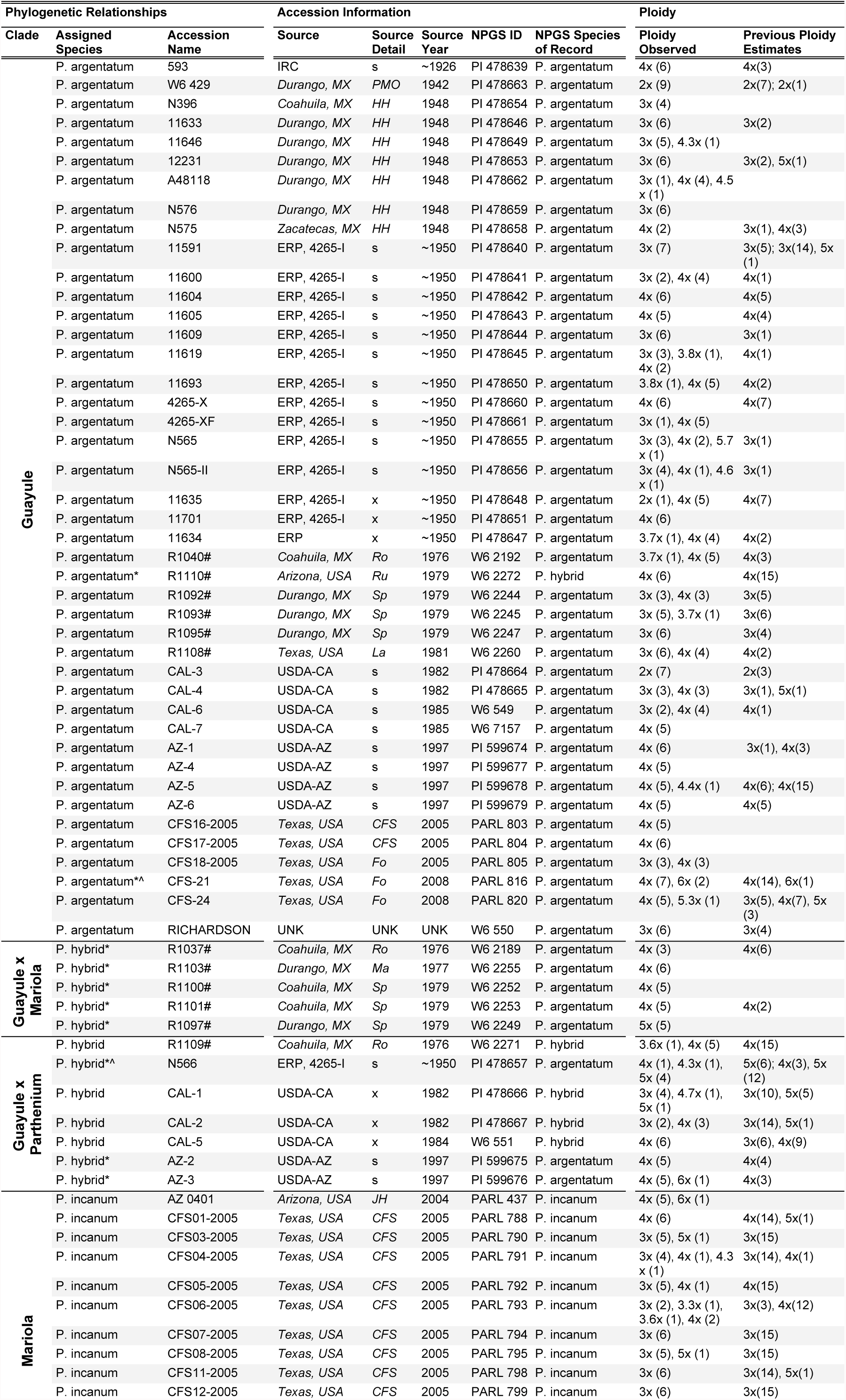
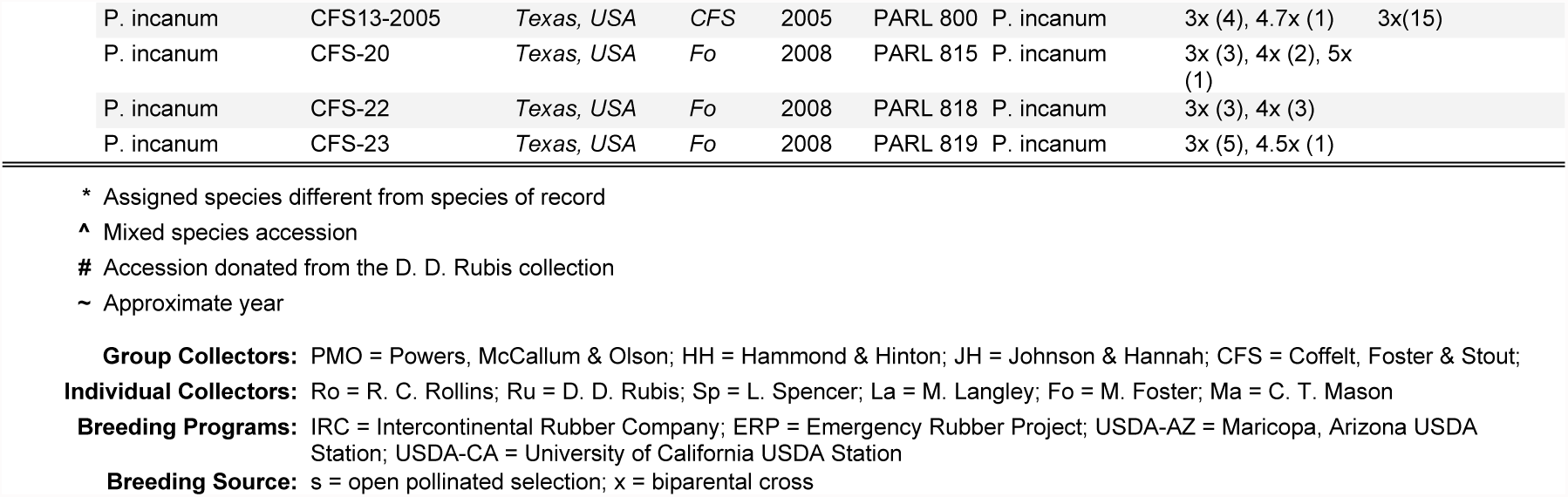
Summary of genotyped accessions from the NALPGRU guayule and mariola germplasm collection. Italic text in the Source and Source Detail columns indicates wild collected accessions. For developed accessions, source information indicates the breeding program followed by development method (s = selection, x = biparental cross). For wild collected accessions, source information indicates the collection location (state, country) followed by collector initials. Ploidy level is denoted by a numeric value indicating the number of chromosome copies (2x = 36 chromosomes) and fractional values indicate aneuploidy. The number of observed samples of a given ploidy level are indicated in parentheses. Multiple sources of previous ploidy estimates are separated by semicolon, and the source publications for those estimates are listed in the references section.

| Phylogenetic Relationships |  |  | Accession Information |  |  |  |  | Ploidy |  |
| --- | --- | --- | --- | --- | --- | --- | --- | --- | --- |
| Clade | Assigned Species | Accession Name | Source | Source Detail | Source Year | NPGS ID | NPGS Species of Record | Ploidy Observed | Previous Ploidy Estimates |
| Guayule | *P. argentatum* | 593 | IRC | s | ~1926 | PI 478639 | *P. argentatum* | 4x (6) | 4x(3) |
|  | *P. argentatum* | W6 429 | *Durango, MX* | *PMO* | 1942 | PI 478663 | *P. argentatum* | 2x (9) | 2x(7); 2x(1) |
|  | *P. argentatum* | N396 | *Coahuila, MX* | *HH* | 1948 | PI 478654 | *P. argentatum* | 3x (4) |  |
|  | *P. argentatum* | 11633 | *Durango, MX* | *HH* | 1948 | PI 478646 | *P. argentatum* | 3x (6) | 3x(2) |
|  | *P. argentatum* | 11646 | *Durango, MX* | *HH* | 1948 | PI 478649 | *P. argentatum* | 3x (5), 4.3x (1) |  |
|  | *P. argentatum* | 12231 | *Durango, MX* | *HH* | 1948 | PI 478653 | *P. argentatum* | 3x (6) | 3x(2), 5x(1) |
|  | *P. argentatum* | A48118 | *Durango, MX* | *HH* | 1948 | PI 478662 | *P. argentatum* | 3x (1), 4x (4), 4.5 x (1) |  |
|  | *P. argentatum* | N576 | *Durango, MX* | *HH* | 1948 | PI 478659 | *P. argentatum* | 3x (6) |  |
|  | *P. argentatum* | N575 | *Zacatecas, MX* | *HH* | 1948 | PI 478658 | *P. argentatum* | 4x (2) | 3x(1), 4x(3) |
|  | *P. argentatum* | 11591 | ERP, 4265-I | s | ~1950 | PI 478640 | *P. argentatum* | 3x (7) | 3x(5); 3x(14), 5x (1) |
|  | *P. argentatum* | 11600 | ERP, 4265-I | s | ~1950 | PI 478641 | *P. argentatum* | 3x (2), 4x (4) | 4x(1) |
|  | *P. argentatum* | 11604 | ERP, 4265-I | s | ~1950 | PI 478642 | *P. argentatum* | 4x (6) | 4x(5) |
|  | *P. argentatum* | 11605 | ERP, 4265-I | s | ~1950 | PI 478643 | *P. argentatum* | 4x (5) | 4x(4) |
|  | *P. argentatum* | 11609 | ERP, 4265-I | s | ~1950 | PI 478644 | *P. argentatum* | 3x (6) | 3x(1) |
|  | *P. argentatum* | 11619 | ERP, 4265-I | s | ~1950 | PI 478645 | *P. argentatum* | 3x (3), 3.8x (1), 4x (2) | 4x(1) |
|  | *P. argentatum* | 11693 | ERP, 4265-I | s | ~1950 | PI 478650 | *P. argentatum* | 3.8x (1), 4x (5) | 4x(2) |
|  | *P. argentatum* | 4265-X | ERP, 4265-I | s | ~1950 | PI 478660 | *P. argentatum* | 4x (6) | 4x(7) |
|  | *P. argentatum* | 4265-XF | ERP, 4265-I | s | ~1950 | PI 478661 | *P. argentatum* | 3x (1), 4x (5) |  |
|  | *P. argentatum* | N565 | ERP, 4265-I | s | ~1950 | PI 478655 | *P. argentatum* | 3x (3), 4x (2), 5.7 x (1) | 3x(1) |
|  | *P. argentatum* | N565-II | ERP, 4265-I | s | ~1950 | PI 478656 | *P. argentatum* | 3x (4), 4x (1), 4.6 x (1) | 3x(1) |
|  | *P. argentatum* | 11635 | ERP, 4265-I | x | ~1950 | PI 478648 | *P. argentatum* | 2x (1), 4x (5) | 4x(7) |
|  | *P. argentatum* | 11701 | ERP, 4265-I | x | ~1950 | PI 478651 | *P. argentatum* | 4x (6) |  |
|  | *P. argentatum* | 11634 | ERP | x | ~1950 | PI 478647 | *P. argentatum* | 3.7x (1), 4x (4) | 4x(2) |
|  | *P. argentatum* | R1040# | *Coahuila, MX* | *Ro* | 1976 | W6 2192 | *P. argentatum* | 3.7x (1), 4x (5) | 4x(3) |
|  | *P. argentatum* * | R1110# | *Arizona, USA* | *Ru* | 1979 | W6 2272 | *P. hybrid* | 4x (6) | 4x(15) |
|  | *P. argentatum* | R1092# | *Durango, MX* | *Sp* | 1979 | W6 2244 | *P. argentatum* | 3x (3), 4x (3) | 3x(5) |
|  | *P. argentatum* | R1093# | *Durango, MX* | *Sp* | 1979 | W6 2245 | *P. argentatum* | 3x (5), 3.7x (1) | 3x(6) |
|  | *P. argentatum* | R1095# | *Durango, MX* | *Sp* | 1979 | W6 2247 | *P. argentatum* | 3x (6) | 3x(4) |
|  | *P. argentatum* | R1108# | *Texas, USA* | *La* | 1981 | W6 2260 | *P. argentatum* | 3x (6), 4x (4) | 4x(2) |
|  | *P. argentatum* | CAL-3 | USDA-CA | s | 1982 | PI 478664 | *P. argentatum* | 2x (7) | 2x(3) |
|  | *P. argentatum* | CAL-4 | USDA-CA | s | 1982 | PI 478665 | *P. argentatum* | 3x (3), 4x (3) | 3x(1), 5x(1) |
|  | *P. argentatum* | CAL-6 | USDA-CA | s | 1985 | W6 549 | *P. argentatum* | 3x (2), 4x (4) | 4x(1) |
|  | *P. argentatum* | CAL-7 | USDA-CA | s | 1985 | W6 7157 | *P. argentatum* | 4x (5) |  |
|  | *P. argentatum* | AZ-1 | USDA-AZ | s | 1997 | PI 599674 | *P. argentatum* | 4x (6) | 3x(1), 4x(3) |
|  | *P. argentatum* | AZ-4 | USDA-AZ | s | 1997 | PI 599677 | *P. argentatum* | 4x (5) |  |
|  | *P. argentatum* | AZ-5 | USDA-AZ | s | 1997 | PI 599678 | *P. argentatum* | 4x (5), 4.4x (1) | 4x(6); 4x(15) |
|  | *P. argentatum* | AZ-6 | USDA-AZ | s | 1997 | PI 599679 | *P. argentatum* | 4x (5) | 4x(5) |
|  | *P. argentatum* | CFS16-2005 | *Texas, USA* | *CFS* | 2005 | PARL 803 | *P. argentatum* | 4x (5) |  |
|  | *P. argentatum* | CFS17-2005 | *Texas, USA* | *CFS* | 2005 | PARL 804 | *P. argentatum* | 4x (6) |  |
|  | *P. argentatum* | CFS18-2005 | *Texas, USA* | *Fo* | 2005 | PARL 805 | *P. argentatum* | 3x (3), 4x (3) |  |
|  | *P. argentatum* **^ | CFS-21 | *Texas, USA* | *Fo* | 2008 | PARL 816 | *P. argentatum* | 4x (7), 6x (2) | 4x(14), 6x(1) |
|  | *P. argentatum* | CFS-24 | *Texas, USA* | *Fo* | 2008 | PARL 820 | *P. argentatum* | 4x (5), 5.3x (1) | 3x(5), 4x(7), 5x (3) |
|  | *P. argentatum* | RICHARDSON | UNK | UNK | UNK | W6 550 | *P. argentatum* | 3x (6) | 3x(4) |
| Guayule x Mariola | *P. hybrid* * | R1037# | *Coahuila, MX* | *Ro* | 1976 | W6 2189 | *P. argentatum* | 4x (3) | 4x(6) |
|  | *P. hybrid* * | R1103# | *Durango, MX* | *Ma* | 1977 | W6 2255 | *P. argentatum* | 4x (6) |  |
|  | *P. hybrid* * | R1100# | *Coahuila, MX* | *Sp* | 1979 | W6 2252 | *P. argentatum* | 4x (5) |  |
|  | *P. hybrid* * | R1101# | *Coahuila, MX* | *Sp* | 1979 | W6 2253 | *P. argentatum* | 4x (5) | 4x(2) |
|  | *P. hybrid* * | R1097# | *Durango, MX* | *Sp* | 1979 | W6 2249 | *P. argentatum* | 5x (5) |  |
| Guayule x Parthenium | *P. hybrid* | R1109# | *Coahuila, MX* | *Ro* | 1976 | W6 2271 | *P. hybrid* | 3.6x (1), 4x (5) | 4x(15) |
|  | *P. hybrid* **^ | N566 | ERP, 4265-I | s | ~1950 | PI 478657 | *P. argentatum* | 4x (1), 4.3x (1), 5x (4) | 5x(6); 4x(3), 5x (12) |
|  | *P. hybrid* | CAL-1 | USDA-CA | x | 1982 | PI 478666 | *P. hybrid* | 3x (4), 4.7x (1), 5x (1) | 3x(10), 5x(5) |
|  | *P. hybrid* | CAL-2 | USDA-CA | x | 1982 | PI 478667 | *P. hybrid* | 3x (2), 4x (3) | 3x(14), 5x(1) |
|  | *P. hybrid* | CAL-5 | USDA-CA | x | 1984 | W6 551 | *P. hybrid* | 4x (6) | 3x(6), 4x(9) |
|  | *P. hybrid* * | AZ-2 | USDA-AZ | s | 1997 | PI 599675 | *P. argentatum* | 4x (5) | 4x(4) |
|  | *P. hybrid* * | AZ-3 | USDA-AZ | s | 1997 | PI 599676 | *P. argentatum* | 4x (5), 6x (1) | 4x(3) |
| Mariola | *P. incanum* | AZ 0401 | *Arizona, USA* | *JH* | 2004 | PARL 437 | *P. incanum* | 4x (5), 6x (1) |  |
|  | *P. incanum* | CFS01-2005 | *Texas, USA* | *CFS* | 2005 | PARL 788 | *P. incanum* | 4x (6) | 4x(14), 5x(1) |
|  | *P. incanum* | CFS03-2005 | *Texas, USA* | *CFS* | 2005 | PARL 790 | *P. incanum* | 3x (5), 5x (1) | 3x(15) |
|  | *P. incanum* | CFS04-2005 | *Texas, USA* | *CFS* | 2005 | PARL 791 | *P. incanum* | 3x (4), 4x (1), 4.3 x (1) | 3x(14), 4x(1) |
|  | *P. incanum* | CFS05-2005 | *Texas, USA* | *CFS* | 2005 | PARL 792 | *P. incanum* | 3x (5), 4x (1) | 4x(15) |
|  | *P. incanum* | CFS06-2005 | *Texas, USA* | *CFS* | 2005 | PARL 793 | *P. incanum* | 3x (2), 3.3x (1), 3.6x (1), 4x (2) | 3x(3), 4x(12) |
|  | *P. incanum* | CFS07-2005 | *Texas, USA* | *CFS* | 2005 | PARL 794 | *P. incanum* | 3x (6) | 3x(15) |
|  | *P. incanum* | CFS08-2005 | *Texas, USA* | *CFS* | 2005 | PARL 795 | *P. incanum* | 3x (5), 5x (1) | 3x(15) |
|  | *P. incanum* | CFS11-2005 | *Texas, USA* | *CFS* | 2005 | PARL 798 | *P. incanum* | 3x (6) | 3x(14), 5x(1) |
|  | *P. incanum* | CFS12-2005 | *Texas, USA* | *CFS* | 2005 | PARL 799 | *P. incanum* | 3x (6) | 3x(15) |

|  |  |  |  |  |  |  |  |  |
| --- | --- | --- | --- | --- | --- | --- | --- | --- |
| P. incanum | CFS13-2005 | Texas, USA | CFS | 2005 | PARL 800 | P. incanum | 3x (4), 4.7x (1) | 3x(15) |
| P. incanum | CFS-20 | Texas, USA | Fo | 2008 | PARL 815 | P. incanum | 3x (3), 4x (2), 5x (1) |  |
| P. incanum | CFS-22 | Texas, USA | Fo | 2008 | PARL 818 | P. incanum | 3x (3), 4x (3) |  |
| P. incanum | CFS-23 | Texas, USA | Fo | 2008 | PARL 819 | P. incanum | 3x (5), 4.5x (1) |  |

\* Assigned species different from species of record
^ Mixed species accession

### Guayule germplasm

The guayule accessions in the NALPGRU germplasm collection (Table 1) are primarily aggregated from four distinct sources: historic guayule accessions derived from the Emergency Rubber Project (ERP) and the 1948 Hammond and Hinton collection (Hammond, 1948), 1970s wild collections of guayule from Mexico obtained from Dr. Rubis’ collection (e.g. R1100, R1037), breeding lines developed during the 1980s and 1990s (accessions AZ-1 through AZ-5 and CAL-1 through CAL-7), and wild collections from Texas, USA in 2005 and 2008 (CFS16-2005, CFS17-2005, CFS18-2005, CF-21, and CFS-24).

Among the 15 ERP-derived accessions, 13 shared as maternal source a phenotypically uniform selection (selection 4265-I; Johnson, 1950) from within a single wild collection (bulk from five plants, collection # 4265; Powers, 1942). The original selection was determined to be primarily triploid (Bergner, 1946; Johnson, 1950). The remaining two ERP-derived accessions are 593, a selection developed by the Intercontinental Rubber Company circa 1926 and transferred to the ERP, and 11634, a selection from an “open-pollinated 36-chromosome cross between SP-7 and SP-8 (N-322 from an old nursery selection S4838-74)” (Hammond and Polhamus, 1965, p. 108). Source history also suggests that three subsequent accessions represent bulk re-capture of guayule genotypes already in the collection: R1110 from abandoned guayule research fields in Mesa, AZ, USA; CFS18-2005 and CFS-24 from abandoned guayule research fields at the Texas Proving Grounds, TX, USA (Ilut et al., 2015)

Based on a PCA of the SNP genotypes, guayule germplasm separates into four major groups (Figure 2) representing triploids recapitulating the 4265-I genotype (group I), hybrids of 4265-I triploids (group II), diverse guayule genotypes from Mexico (group III), and novel guayule genotypes from Big Bend National Park, Texas, USA (group IV). Group I includes four accessions derived from 4265-I (11591, 11619, N565, N565-II), two Hammond and Hinton accessions (11646 and N576), the RICHARDSON accession, two triploids from the CAL-6 accession, and one triploid from CFS18-2005. Both the PCA (Figure 2) and the phylogenetic tree (Figure 3) indicate that all of these triploid samples are very closely related and likely share a common origin, while historical information suggests these genotypes are representative of the 4265-I accession developed by Johnson in 1950. Provenance information is inconclusive at best for most of these accessions. Accessions 11646 and N576 pre-date the 1950 selection that created 4265-I, but it would appear they were collected from the same general area (southeast Durango, Mexico; Hammond and Polhamus, 1965, p. 105) as the original 4265 accession. The source of accession CAL-6 is a triploid selection from guayule genotypes present in the germplasm collection prior to 1981 (Estilai, 1986). It currently contains a mixture of tetraploid and triploid genotypes, but the source accession for the original selection is unknown. The CFS18-2005 accession represents a bulk re-capture of guayule genotypes present in the germplasm collection prior to 1978, in agreement with the observation that samples from this accession are found in groups I, II, and III. Finally, there is no information available regarding the collection locale or accession source of the bulk RICHARDSON accession.

**Figure 2.**
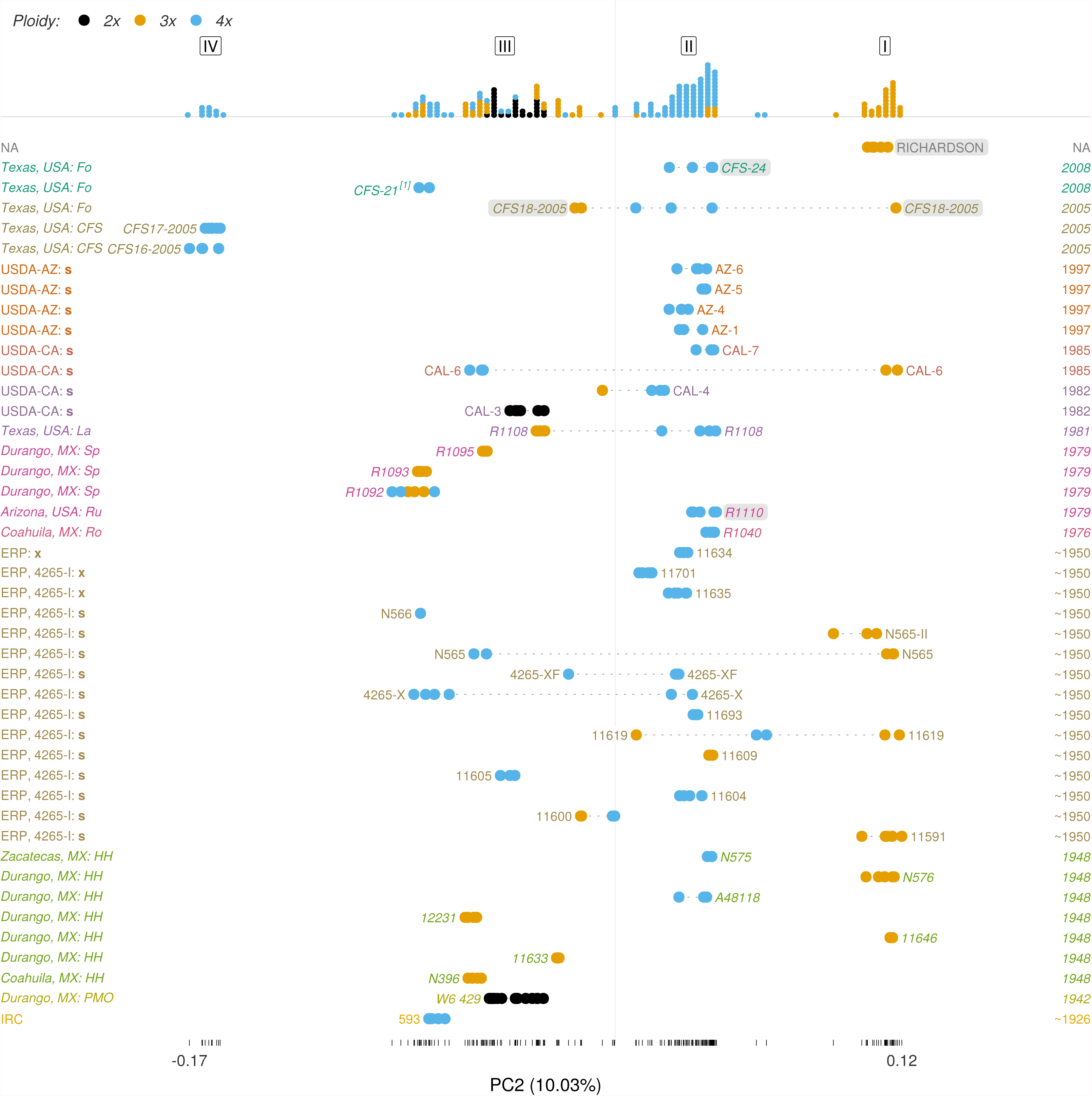
Genetic diversity of guayule accessions. Italic text indicates wild collected accessions, and point colours indicate the ploidy level of individual samples (black = diploid, orange = triploid, blue = tetraploid). The x axis represents a major dimension of genetic diversity (PC2, explaining 10% of the observed genetic diversity) concordant with the phylogenetic tree in Figure 3, and the axis line is marked with a vertical dash at each observed value. The y axis represents individual accessions ordered chronologically by the year in which they were collected or created (increasing, bottom to top). The year is indicated on the right axis, and the source is indicated on the left axis.

In group II, there is a broad selection of accessions, almost exclusively tetraploid. Phylogenetically they are relatively diverse (Figure 3), but the PCA (Figure 2) indicates they are more closely related to group I than all other accessions in the collection. Given their tight clustering equidistant from group I, and the historical information available on accessions in this group, they are most likely the result of hybridization (and, in most cases, ploidy increase) between triploid 4265-I plants and other guayule genotypes. Among ERP-derived accessions, six of the seven historical tetraploid open-pollinated selections from (or crosses with) triploid accession 4265-I are either entirely (11604, 11693, 11635, 11701) or partially (4265-X, 4265-XF) included in this group. Accession 11634 likewise belongs to this group, suggesting that this tetraploid genotype was derived from the original diploid parent by open-pollinated outcrossing with the ubiquitous 4265-I genotype. In addition to the tetraploids, the placement of triploid accession 11609 within this group suggests that this accession is derived from a sexual outcrossing of triploid 4265-I plants.

**Figure 3.**
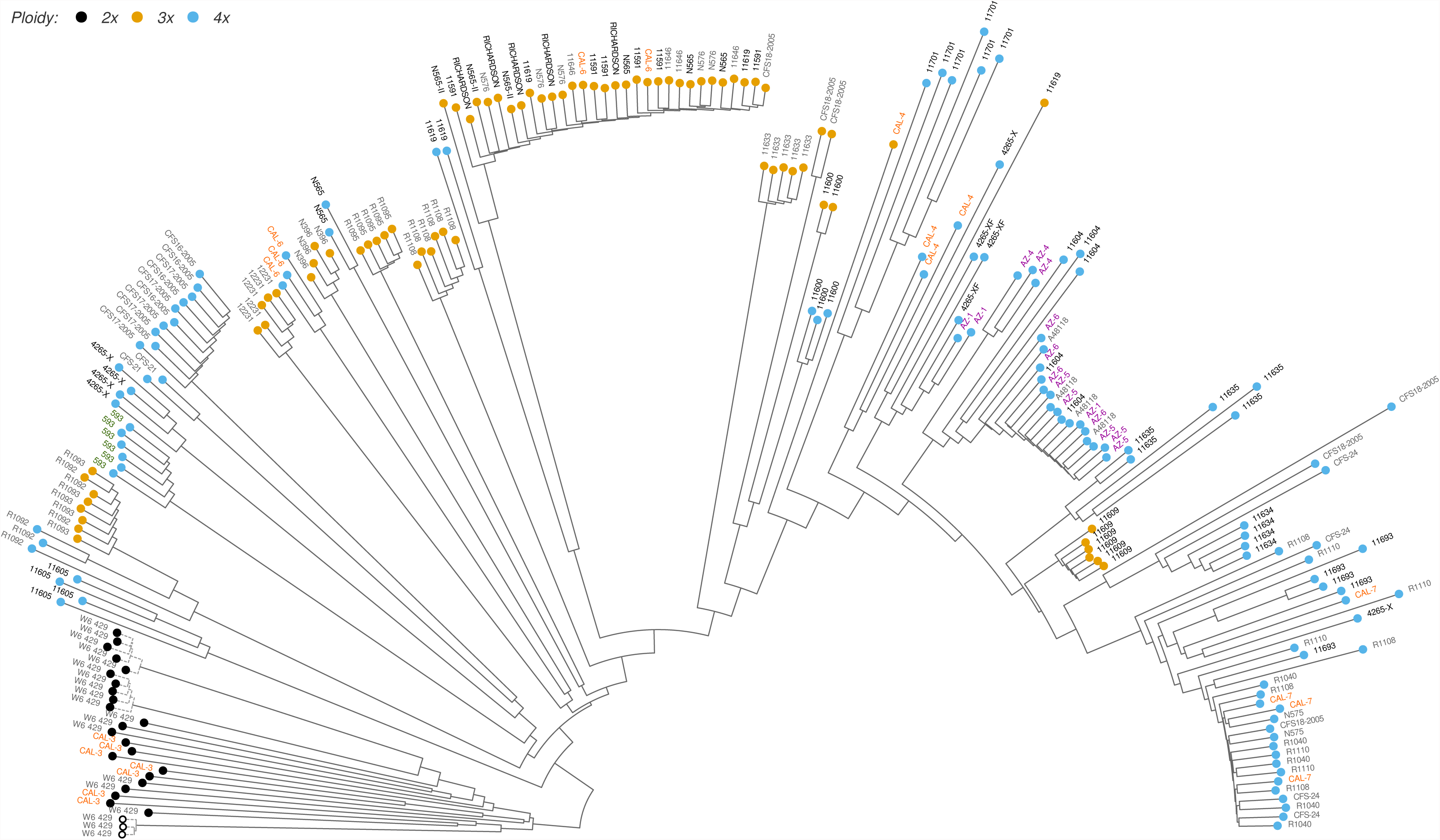
Phylogenetic tree of guayule accessions. For visualization purposes, the phylogenetic tree is rooted between diploid and polyploid guayule samples. Samples are labelled with accession names (gray text = wild collected accessions). Specific breeding programs are colour coded as follows (abbreviations as in Table 1): IRC = green; ERP = black; USDA-AZ = purple; USDA-CA = orange. Terminal node colours indicate the ploidy level of individual samples (black = diploid, orange = triploid, blue = tetraploid). The samples used to create the de-novo GBS reference are marked by open circle terminal nodes. Dashed branches represent technical replicates of the same plant.

The two Hammond and Hinton collections in group II were once again collected from the same general area as the original 4265 collection. Accession A48118 was collected from the same area in southeast Durango as N576 (an accession in group I), and accession N575 “was collected nearby in northern Zacatecas” (Hammond and Polhamus, 1965, p. 105). Other wild collections in this group include tetraploids from bulk re-capture accessions (R1110, CFS18-2005, CFS-24), accession R1108, and accession R1040. Although it is not surprising to find this genotype group represented in bulk re-capture accessions, historical collection data do not provide plausible explanations for its presence in R1108 (a 1981 wild collection from Big Bend, Texas, USA) and R1040 (ostensibly a 1976 wild collection from Ocampo, Coahuila, Mexico). Phylogenetic analysis (Figure 3) indicates that tetraploid samples from these two accessions are very closely related to the two samples from wild collection N575 as well as subsequent bulk re-capture (CFS24, CFS18-2005) or anonymous source selection (CAL7) accessions, and the entire group is most closely related to accession 11693. In contrast, triploid samples from R1108 form a distinct, unique phylogenetic group, as would be expected based on their collection date and locale. The most parsimonious explanation for this discrepancy would be a historical contamination of seed collections from accessions R1108 and R1040 with seed from accession N575 or 11693, and further genotyping of samples from these two accessions is recommended in order to elucidate this issue.

The remaining accessions in group II (CAL-4, CAL-7, AZ-1, AZ-4, AZ-5, AZ-6) represent selections from either other accessions in this group (AZ-1, derived from 4265-X; Ray et al., 1999) or anonymous accessions present in the germplasm collection prior to 1981 (CAL-7, Estilai, 1986; AZ-4, AZ-5, AZ-6, Ray et al., 1999; CAL-3, Tysdal et al., 1983). Phylogenetic analysis of the SNP marker data indicates that AZ-5 and AZ-6 contain genotypes derived from 11604 and A48118, with AZ-4 distinct from but most closely related to all four of these accessions (Figure 3). AZ-1 is most closely related to samples from 4265-X and 4265-XF(a selection from 4265-X), as expected. CAL-4, a composite open-pollinated seed collection, is composed of genotypes from multiple accessions, including several samples most closely related to 4265-XF. As noted above, the historical source for CAL-7 is unknown, and the possibility of seed contamination in the accessions to which it is most closely related makes it difficult to assign a putative source accession based on SNP marker data.

In contrast with the previous two groups, group III contains a mixture of ploidy levels (diploid, triploid, tetraploid), is mainly composed of wild collected guayule from Mexico, and accessions within this group tend to form distinct, independent clades (Figure 2, Figure 3). This group includes the oldest guayule accession (593, a tetraploid accession selected from Mexican germplasm circa 1926), the diploid guayule samples (W6 429, originally collected near Mapimí, Durango, Mexico in 1942, and CAL-3, selected in 1982 from W6 429), triploid wild collections from Coahuila, Mexico in 1948 and Durango, Mexico in 1948 and 1971, triploid wild collections from Big Bend, Texas, USA in 1981, and tetraploid collections from Bakerfield, Texas, USA in 2008. In addition to wild collections, ERP-derived accessions are represented in this group by accessions 11600 and 11605 as well as several samples from 4265-X, 4265-XF, N565, and N566. It is unclear at this point whether samples from these ERP-derived accessions or tetraploid samples from accession CAL-6 (originally a triploid selection of pre-1981 germplasm) represent contamination of seed from 4265-I derived accessions with seed from wild collections or further outcrossing of these accessions during open-pollinated seed increases. Accessions from this diverse, genetically heterogeneous group are ideal candidates for exploring guayule genetic diversity in future breeding programs.

The final group, group IV, consists of tetraploids from two accessions collected in 2005 along the same road in Big Bend National Park, Texas, USA. These two accessions are indistinguishable from each other based on SNP markers (Figure 3), and they represent the most diverged genotype group when compared to the rest of the collection. This suggests that future collections of wild guayule from within Big Bend National Park are likely to add novel genotypes to the collection and increase the genetic diversity present therein.

Complementary to phylogenetic analysis across accessions, an analysis of heterozygosity within samples provides a measure of genetic diversity that can inform our understanding of ploidy variation mechanisms. Given that all SNP genotyping was performed against a diploid reference, exact heterozygosity values are generally correlated with the ploidy level of a given sample (Supplemental Figure A9). However, tetraploid guayule samples exhibit a broad range of heterozygosity levels, which suggests that they are the result of both self-fertilization and outcrossing. Although polyploid guayule is generally apomictic, fertilized embryos are occasionally produced (see Thompson and Ray, 1989, p. 106 for a review of guayule reproduction) and self-fertilization can lead to a marked reduction in observed heterozygosity in the progeny. The data suggest that self-fertilization accompanied by reduction of the megaspore mother cell (Class IV in Thompson and Ray, 1989 Table 4.1) is relatively common among tetraploid guayule.

### Guayule hybrids and mariola

Guayule and mariola are known to readily form hybrids both in the wild (Rollins, 1944) and under experimental conditions (Powers and Rollins, 1945). Moreover, historically, morphological characteristics of these hybrids have often been ascribed to guayule (Rollins, 1950). It is therefore not surprising that several guayule accessions in the germplasm collection were identified as guayule-mariola hybrids. In particular, the genotyping data indicates that five accessions collected in the 1970s from Mexico (R1037, R1097, R1100, R1101, R1103) are guayule-mariola hybrids (Figure 1; Figure 4). Phylogenetic analysis and PCA indicate that pentaploid hybrid accession R1097 is a hybrid of guayule triploid accession R11095 and an unknown mariola genotype, and hybrid accession R1103 is derived from a guayule parent with a genotype distinct from those in the current guayule germplasm collection (Figure 4, PC2).

**Figure 4.**
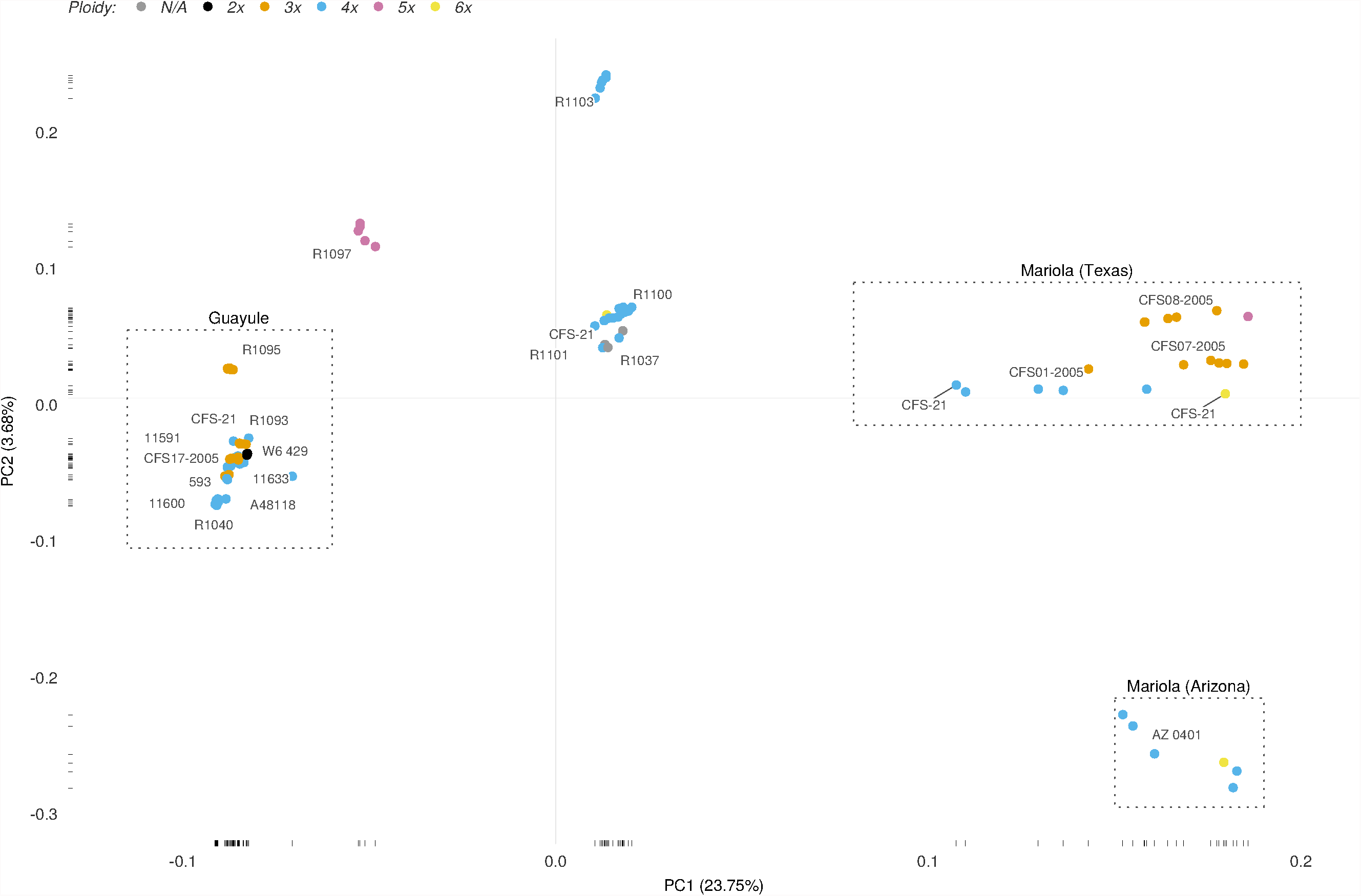
Genetic diversity of guayule-mariola hybrids. Only the top two principal components are shown, and the proportion of variation explained by each component is indicated on the axis labels. Outgroup samples are demarcated by dashed boxes and labelled above each box. Representative accessions for outgroups of guayule and mariola were selected based on genetic diversity within those clades. Accession labels are placed at the median coordinates for samples within those accessions and slightly offset for legibility. Labels attached to individual data points are indicated by a line connecting the label and point. For accession CFS-21, which contains both outgroups and hybrids, samples were labelled separately within each group. Point colours indicate the ploidy level of individual samples: gray = unknown ploidy, black = diploid, orange = triploid, blue = tetraploid, purple = pentaploid, yellow = hexaploid.

Early experiments (Rollins, 1946; Tysdal, 1950) indicated that guayule was also amenable to hybridization with other *Parthenium* species besides mariola, and three interspecific crosses were created in 1978 (Waln et al., 1983) and released as new accessions in 1982 and 1984 as CAL-1 (*Parthenium argentatum* Gray x *Parthenium tomentosum* var. *tomentosum* DC), CAL-2 (*Parthenium argentatum* Gray x *Parthenium fruticosum* Less.), and CAL-5 (*Parthenium argentatum* Gray x *Parthenium tomentosum* var. *stramonium* (Green) Rollins) by the University of California, Davis and the USDA Cotton Research Station in Shafter, California, USA (Estilai, 1985; Tysdal et al., 1983). In addition to these known non-mariola hybrids, phylogenetic analysis and PCA both indicate that wild collected accession R1109, accessions AZ-2 and AZ-3, and pentaploid samples from accession N566 are also hybrids with non-mariola *Parthenium* species (Figure 1; Figure 5). PCA suggests that the non-guayule parents of these four hybrids are one (or several closely related) species distinct from *P*. *tomentosum* and *P*. *fruticosum* (Figure 5, PC1). Moreover, the guayule parent of the hybrids R1109, AZ-2, and AZ-3 is likely not sampled in the existing germplasm pool (Figure 5, PC2).

**Figure 5.**
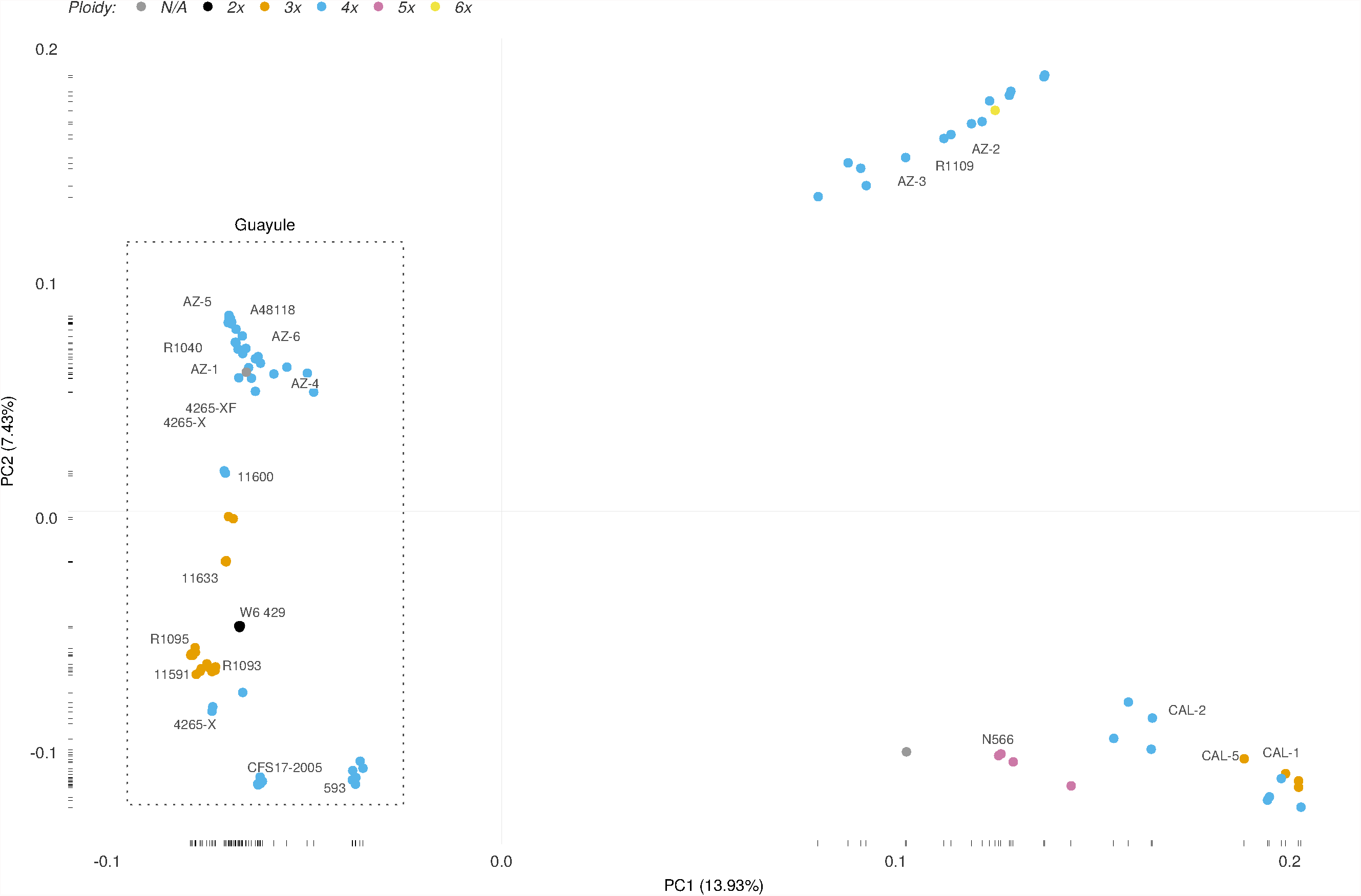
Genetic diversity of non-mariola guayule hybrids. Only the top two principal components are shown, and the proportion of variation explained by each component is indicated on the axis labels. Guayule outgroup samples are demarcated by a dashed box. Representative outgroup accessions of guayule were selected based on genetic diversity within the guayule clade. Accession labels are placed at the median coordinates for samples within those accessions and slightly offset for legibility. Point colours indicate the ploidy level of individual samples: gray = unknown ploidy, black = diploid, orange = triploid, blue = tetraploid, purple = pentaploid, yellow = hexaploid.

In addition to guayule and hybrid accessions, we analyzed 14 accessions of mariola: one collected from Arizona, USA (accession AZ 0401) and 13 collected from southwestern Texas, USA in between 2004 and 2008 (Supplementary Table A2). The mariola genotypes sampled in this study are relatively homogeneous, with the PCs explaining at most 3% of the observed variance (Supplementary Figure A4).

### Mixed and conflicting accessions

Overall, the observed genetic relationships between accessions are in agreement with expected relationships based on collection and breeding history. Among guayule accessions, eight accessions exhibit larger than expected genotypic diversity (Figure 2, accessions with horizontal dashed lines), suggesting a mixture of genotypes within each accession. The ERP-derived accessions 11619, 4265-X, 4265-XF, and N565 contain, in addition to the expected genotypes from groups I and II (Figure 2), multiple samples from the diverse group III, suggesting historical contamination of the seed source for these accessions with seed from other contemporary accessions. Accession R1108, collected from Big Bend, Texas, USA in 1981, contains two distinct genetic groups: a unique triploid genotype, and a tetraploid genotype from group II. These relationships suggest that the triploid genotypes in R1108 represent the original wild collection while the tetraploid genotypes are an indication of subsequent hybridization with or seed contamination from ERP-derived accessions. Accessions CAL-4 and CAL-6 are selections from anonymous, pooled germplasm (Estilai, 1986), and accession CFS18-2005 represents a bulk collection of feral guayule cultivars from an abandoned guayule research site at the Firestone Proving Grounds, Texas, USA (Ilut et al., 2015). It is therefore not surprising to find that they contain a broad range of genotypes similar to existing genotypes in the germplasm collection.

Two accessions (N566 and CFS-21) contain samples from multiple species or hybrids (Table 1; Supplementary Table A1). Accession N566 contains a mixture of tetraploid and pentaploid samples, with the tetraploids similar to historic accession 593 and pentaploids hybrids between guayule and other non-mariola *Parthenium* species. Previous studies (Gore et al., 2011) have also observed a mixture of tetraploids and pentaploids in this accession. Based on historical information, this accession is expected to be an open-pollinated selection from 4265-I. However, genetic relationships between guayule and guayule hybrids (Figure 5, PC1) indicate that the guayule parent of these hybrids is more similar to accession 593 rather than accessions such as 11591 (a representative accession for the historic 4265-I genotype). Given that accession 593 precedes the ERP and was used extensively during that program, it is plausible that accession N566 represents an interspecific hybrid between guayule accession 593 and an unknown *Parthenium* species. The other multi-species accession (CFS-21) was collected from Texas, USA in 2008 and contains samples of guayule, mariola, and guayule-mariola hybrids. However, it would appear that the current source of seed for this accession contains a mixture of guayule and mariola plants, and the seed stock center is currently in the process of re-planting this accession from original seed (Heinitz, 2017). Given that the guayule samples in this accession represent unique genotypes, it would be advisable to re-evaluate this accession in the future as a possible source of novel guayule genotypes.

The marker-based analysis indicates that several accessions should be reclassified (Table 1). Accession R1110, collected from Mesa, Arizona, USA in 1979, is currently classified as a hybrid. However, genotype data indicates that this accession is indeed guayule, and belongs to guayule group II (4265-I derived accessions). Given that this accession was collected from a previous guayule research station (Ilut et al., 2015), it is likely that it represents a re-capture of existing, ERP-derived genotypes. Five other accessions (R1037, R1097, R1100, R1101, R1103) collected during the 1970s from Mexico and currently classified as guayule are guayule-mariola hybrids. A sixth, R1109, also currently classified as guayule, is a hybrid between guayule and an unknown non-mariola *Parthenium* species.

Finally, two of the most commonly used contemporary accessions, AZ-2 and AZ-3, are also hybrids between guayule and an unknown non-mariola *Parthenium* species. Although the recorded source accession for these selections is 11591 (Ray et al., 1999), there is some uncertainty as to the exact source and they are the result of multiple rounds of open-pollinated selection (Dierig, 1987; Ray, 2014). The original plant material for these selections came from the same field as accession R1109 (Ilut et al., 2015), and, given their high genetic similarity to accession R1109, it is therefore plausible that these accessions were derived from it or hybridized with the same *Parthenium* species.

It should be noted that this list of potentially problematic accessions is by no means exhaustive. In order to focus the primary analysis on the genotypes most likely to be representative of a given accession, discordant, single sample genotypes were removed. Phylogenetic analysis results that include all samples for a given accession are presented in Supplementary Figures A5-A8, and they indicate that many accessions are likely to contain minority genotypes from sources other than the accession origin. It is therefore imperative that plants used in future experiments or breeding programs are genotyped individually rather than relying on accession information for their likely genotype.

## Conclusions

The existing germplasm collection represents primarily wild-collected material or selections from such material. However, based on historical information, we expected a large proportion of accessions to have very similar genotypes derived from a single triploid source (4265-I). Several accessions (Figure 2, group I) do indeed recapitulate that genotype, but tetraploid selections from that source (Figure 2, group II) are most likely the result of outcrossing fertilization (Class II in Thompson and Ray, 1989 Table 4.1) and thus capture some of the genetic diversity present in the hundreds of ERP accessions that are no longer available. In addition, several wild collections from Mexico in the 1940s and 1970s (Figure 2, group III) provide hitherto untapped diversity for guayule breeding. Furthermore, the genetically distinct accessions collected in Big Bend National Park, Texas, USA in 2005 (CFS16-2005, CFS17-2005; Figure 2, group IV) indicate that future collections from this location have tremendous potential to expand available genetic diversity. Overall, genetic diversity in the guayule germplasm collection is higher than expected based on historical information.

In addition to characterizing genetic diversity, we identified several interspecific hybrid accessions of guayule. Although guayule is the only *Parthenium* species that produces commercially exploitable amounts of natural rubber, interspecific hybrids of guayule have the potential to greatly improve the agronomic traits of the plant. There is concern, however, that such interspecific hybrids would result in unacceptable decrease in the amount of natural rubber produced. It is therefore encouraging to note that two of the most commonly used high yielding accessions (AZ-2 and AZ-3) are interspecific hybrid guayule accessions. This indicates that guayule interspecific hybrids provide a viable path for agronomic improvement of this nascent commercial crop.

This study provides the most thorough genetic characterization of guayule germplasm to date, and provides the necessary foundation for the development and rapid improvement of commercial guayule germplasm using genomics-assisted breeding strategies. Due to the apomictic nature of polyploid guayule reproduction, standard breeding approaches such as biparental crosses still remain a challenge. However, the genotyping methods described here and the genetic diversity characterized provide the tools to explore, dissect, and hopefully control the degree of apomixis in polyploid guayule.

## Data Availability

All sequence data have been deposited under BioProjectPRJNAxxxxxx in the NCBI Sequence Read Archive (SRA; Wheeler et al., 2008). For each plant sample, the corresponding SRA accession number is indicated in Table X.X (see Supplementary data). Nucleotide sequences for all the de-novo assembled reference loci, unfiltered genotype calls, and locus sequences and SNPs for the high confidence SNP data set are provided as an online supplemental data set.

## Materials and Methods

### Plant material, growth conditions, and ploidy analysis

Seeds (achenes) for 69 accessions were obtained from the USDA-ARS National Plant Germplasm System (NPGS; www.ars-grin.gov/npgs) through the National Arid Land Plant Genetics Resources Unit (NALPGRU) in Parlier, CA. Plants were greenhouse grown as described (Sanchez et al., 2014). Leaf tissue samples were collected and prepared for ploidy analysis as previously described (Ilut et al., 2015, sec. 2.3; Sanchez et al., 2014), with an average of two technical replicates for each plant. For each plant, we calculated its ploidy level as described in Sanchez et al. (2014), then assigned an integer-valued ploidy level if the nearest integer value was within two standard deviations of the mean across technical replicates, and a fractional value (denoting likely aneuploid samples) otherwise. Supplementary Table A1 contains detailed source information for all the plants genotyped in this study, including NALPGRU seed inventory identifiers and detailed ploidy measurements.

### DNA extraction, sequencing, and genotyping

On average, we selected six plants (samples) per accession and followed the protocols of Ilut et al. (2015) for tissue sampling, DNA library construction, and sequencing (Ilut et al., 2015, sec. 2.6), read filtering (Ilut et al., 2015, sec. 2.7), and de-novo GBS reference sequence construction (Ilut et al., 2015, sec. 2.8). We selected samples 72_1_1-15, 72_1_1-l3, and 72_1 (see Supplementary Table A1), which represent three technical replicates of the same tissue source, for the construction of a de-novo GBS reference sequence.

The trimmed and filtered reads for each sequencing sample were aligned against the de-novo GBS reference sequence using bwa-mem (version 0.7.12-r1039; Li, 2013), and GATK (version 3.5; McKenna et al., 2010) was used to perform indel realignment. The HaplotypeCaller and CombineGVCFs modules of GATK were used to generate sample-level genotype calls and produce combined genotyping results for all samples.

### Estimation of heterozygosity levels

Heterozygosity was estimated independently for each sample. For a given sample, we extracted all nucleotides in the de-novo reference which had at least 10 reads aligned to them with a mapping quality of 30 or better using samtools (version 1.2-25; Li et al., 2009) and calculated the proportion of these nucleotides with heterozygous genotype calls. Prior to calculating this proportion, the genotype data were filtered using vcftools (version 0.1.14; Danecek et al., 2011) such that all genotype calls with genotype quality below 30 were marked as missing and any SNPs with SNP quality below 30 were removed. Heterozygosity estimates were not adjusted for ploidy level.

### Phylogenetic analysis

For phylogenetic analysis, genotype data were filtered using vcftools in a three stage process. First, only biallelic SNPs with a SNP quality of 30 or better were retained, and genotype calls with a read depth less than 4 or a genotype quality less than 30 were set to missing. Second, any SNPs with more than 20% missing data were removed. Third, any samples with more than 50% missing data were removed. This filtering was performed independently on the four major groups described in Figure 1 (Guayule, *Parthenium* Hybrids, Mariola Hybrids, Guayule).

Phylogenetic networks and neighbour joining trees were constructed using SplitsTree (version 4.14.4; Huson and Bryant, 2006), and nodes with bootstrap support of less than 75% over 10,000 replicates were collapsed. Principal component analysis (PCA) was performed using the smartpca component of EIGENSOFT (version 6.1.4; Patterson et al., 2006). Aneuploid samples were removed from the primary phylogenetic analysis, and samples within each of the four major groups (Figure 1) were filtered further based on PCA results by removing any samples within an accession that were genotype outliers. Genotype outliers were defined as outliers within an accession along one or more PCs such that the sum of the variance explained by these PCs was 10% or more. Outliers were identified using the quantile method as implemented in the stats package of the R statistical computing environment (version 3.2.2; R Core Team, 2015), with a maximum quartile distance threshold of three interquartile ranges.

## Acknowledgements

We thank the National Arid Land Plant Genetics Resources Unit at Parlier, CA, USA for providing seeds. Also, we wish to thank Greg Leake and Brenda Singleton for maintaining plants, and Nicholas LaForest for leaf sample collection and ploidy analysis at the USDA-ARS, U.S. Arid-Land Agricultural Research Center. Preparation of total genomic DNA samples for GBS was carried out by Nicholas Kaczmar in the Gore laboratory at Cornell University. We thank Olivia Tonge for assistance in a preliminary phylogenetic analysis of diploid guayule samples. This work was funded and supported by the USDA-ARS and USDA-NIFA/DOE Biomass Research and Development Initiative (BRDI) Grant No. 2012-10006. Mention of trade names or commercial products in this publication is solely for the purpose of providing specific information and does not imply recommendation or endorsement by the United States Department of Agriculture. The USDA is an equal opportunity provider and employer.

## Supplemental Files

Table A1. Detailed sample summary

Table A2. Detailed accession summary

Table A3. SNP selection for phylogenetic analysis

Figure A1. Detailed PCA of selected samples of guayule

Figure A2. Detailed PCA of selected samples of non-mariola guayule hybrids

Figure A3. Detailed PCA of selected samples of mariola guayule hybrids

Figure A4. Detailed PCA of selected samples of mariola

Figure A5. Detailed PCA of all samples of guayule. Outlier samples are indicated by open circles. Aneuploid samples are indicated by open circles and black outer circle.

Figure A6. Detailed PCA of all samples of non-mariola guayule hybrids. Outlier samples are indicated by open circles. Aneuploid samples are indicated by open circles and black outer circle.

Figure A7. Detailed PCA of all samples of mariola guayule hybrids. Outlier samples are indicated by open circles. Aneuploid samples are indicated by open circles and black outer circle.

Figure A8. Detailed PCA of all samples of mariola. Outlier samples are indicated by open circles. Aneuploid samples are indicated by open circles and black outer circle.

Figure A9. Heterozygosity distribution for select samples guayule, guayule hybrids, and mariola

Figure A10. Heterozygosity distribution for all samples guayule, guayule hybrids, and mariola. Aneuploid samples are indicated by a black outer circle.

